## Supplementary Text and Figures for "Single-cell dynamics of pannexin-1-facilitated programmed ATP loss during apoptosis"

Supplementary information

### Supplementary Methods

#### Chemicals

Probenecid was obtained from Sigma-Aldrich. Solution of probenecid was neutralized with sodium hydroxide before use. Recombinant human Trail and anti-His tag antibody were purchased from R&D. Trail was dissolved in DPBS(-) supplemented with 0.1% BSA at a concentration of 20 μg/mL, and then stored at -80˚C. Anti-His antibody was dissolved in DPBS(-) at a concentration of 1 mg/mL, and then stored at -80˚C. Alexa647-annexin-V were purchased from Molecular Probes.

#### Mammalian cell culture and gene knock out

The SW480 cell line was obtained from American Type Culture Collection. The cells were cultured in Leibovitz's L-15 medium (Wako Pure Chemicals, Japan) supplemented with 10% FBS. Apoptosis of SW480 cells was initiated by adding Trail (50 ng/mL) and anti-His tag antibody (1 μg/mL). Knockout of PANX1 gene was carried out using pre-designed PANX1-KO CRISPR-Cas9 plasmids (Santa Cruz Biotechnology). Knockout of PANX1 gene was verified by western blotting and by sequencing of the targeted region of the genomic DNA.

#### Western blotting

Cleavage of ATeam in apoptotic cells was examined by western blotting. Briefly, apoptosis was induced in HeLa cells, which were previously transfected with pcDNA-AT1.03 or pcDNA-AT1.03CR, by adding anti-FAS antibody and CHX. The cells were harvested 12 h after anti-FAS stimulation and lysed. After separation of the cell lysate by SDS-PAGE, ATeam and actin proteins were detected by western blotting using a polyclonal anti-GFP antibody (Invitrogen), which cross-reacts with both CFP and YFP, and a monoclonal anti-actin antibody (Millipore), respectively. HRP-labeled anti-rabbit and anti-mouse IgG antibodies (GE Healthcare) were used as secondary antibodies.

#### Fluorescence imaging of caspase-3 activity and $\Delta\Psi_m$ during apoptosis

HeLa cells were transfected with the pcDNA-SCAT3.1 plasmid (Nagai and Miyawaki, 2004) using Lipofectamine 2000 (Thermo Scientific). One day after transfection, cells were trypsinized and plated on a collagen-coated glass-bottom dish (0.16 – 0.19 mm thick; MatTek). Two days after transfection, the medium was replaced by phenol red-free DMEM containing 10% FBS and 50 nM TMRE. Then, the cells were visualized with a Ti-E inverted microscope (Nikon, Tokyo, Japan) using a Plan Apo 40×, 0.95 numerical aperture, dry objective lens (Nikon). Cells were maintained on a microscope at 37 °C with a continuous supply of a 95% air and 5% carbon dioxide mixture by using a stage-top incubator (Tokai Hit). All filters used for fluorescence imaging were purchased from Semrock (Rochester, NY): for dual-emission ratio imaging of SCAT3.1 biosensors, an FF01-438/24 excitation filter, an FF458-Di02 dichroic mirror, and two emission filters (an FF02-483/32 for CFP and an FF01-542/27 for YFP); for imaging of TMRE, an FF01-562/40 excitation filter, an FF593-Di02 dichroic mirror, and an FF01-641/75 emission filter. Cells were illuminated using a 75 W xenon lamp through 25% and 12.5% neutral density filters. Fluorescence emissions from cells were imaged using a Zyla4.2 scientific CMOS camera (Andor Technologies). The microscope system was controlled by NIS-Elements software (Nikon). Image analysis was performed using MetaMorph software (Molecular Devices).

**Fluorescence imaging of phosphatidylserine**

HeLa cells were cultured on a 35 mm glass-bottom culture dish. At 6 hours before imaging, the medium was replaced by 2 mL of phenol red-free DMEM supplemented with FBS (10%), $CaCl_2$ (1 mM), propidium iodide (1 μg/mL), annexin V-Alexa647 (10 μL), anti-FAS (250 ng/mL) and CHX (10 μM). Cells were visualized with a Ti-E inverted microscope (Nikon, Tokyo, Japan) using a Plan Apo 20×, 0.75 numerical aperture, dry objective lens (Nikon). All filters used for fluorescence imaging were purchased from Semrock (Rochester, NY): for imaging of propidium iodide, an FF01-504/12 excitation filter, an FF593-Di02 dichroic mirror, and an FF01-562/40 emission filter; for imaging of annexin V-Alexa647, an FF02-628/40 excitation filter, an FF660-Di02 dichroic mirror, and an FF01-692/40 emission filter. Fluorescence emissions from cells were imaged using a Zyla4.2 scientific CMOS camera (Andor Technologies). Image analysis was performed using MetaMorph software (Molecular Devices). Fluorescence intensity of Alexa647 within a whole-cell area of each cell that was shrunk but did not exhibit propidium iodide fluorescence was quantified.

**Fluorescence imaging of free $Mg^{2+}$**

$Mg^{2+}$ imaging was performed using a synthetic $Mg^{2+}$ indicator MGH (Matsui et al., 2017). HeLa cells maintained in 10% FBS in DMEM (Invitrogen) at 37 ºC under 5% $CO_2$ were transfected with the plasmid pcDNA-3.1-(+)-HaloTag using PEI-Max (Polysciences). After 48 h, the cells were washed twice with HBSS and incubated with 5 μM MGH(AM) for 30 min, then 50 nM Halo-TMR for 100 min. After washing the cells twice with HBSS, the medium was replaced by phenol red-free DMEM containing 10% FBS. After 4 h, the cells were washed with HBSS. Imaging of the cells was started just after replacing the medium by phenol red-free DMEM containing 10% FBS, 50 ng/mL anti-FAS and 10 μM cycloheximide. Cells were visualized with a Ti-E inverted microscope using a Plan Apo 40×, 0.95 numerical aperture, dry objective lens (Nikon). Cells were maintained on a microscope at 37 °C with a continuous supply of a 95% air and 5% carbon dioxide mixture by using a stage-top incubator (Tokai Hit). For imaging of MGH, an FF01-497/16 excitation filter, an FF516-Di01 dichroic mirror and an FF01-535/22 emission filter were used. For imaging of TMR, an FF01-562/40 excitation filter, an FF593-Di02 dichroic mirror, and an FF01-641/75 emission filter were used. All filters were purchased from Semrock.

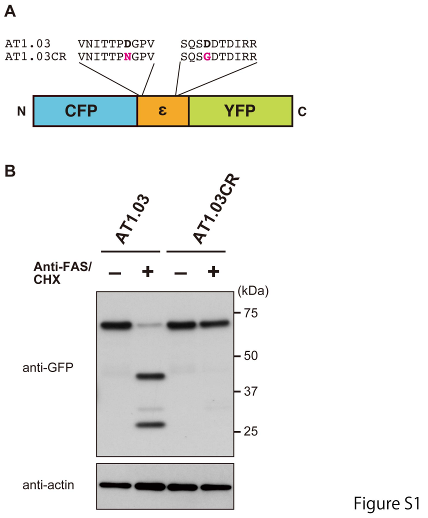


**Figure S1. AT1.03CR, a caspase-resistant mutant of the FRET-based ATP biosensor ATeam.** (A) Schematic drawing of ATeam. An ATP binding protein (*Bacillus subtilis* $F_oF_1$-ATP synthase ε subunit) is sandwiched by CFP (mseCFP) and YFP (cp173-mVenus). Critical Asp residues (Asp-242 and Asp-339) within putative caspase-cleavage sequences within the original ATeam (AT1.03) were replaced by Asn and Gly, respectively. (B) Western blot analysis of the integrity of ATeam in apoptotic cells revealed that AT1.03CR was resistance to cleavage upon the initiation of apoptosis. HeLa cells expressing either AT1.03 or AT1.03CR were stimulated with anti-FAS and cycloheximide (CHX) to induce apoptosis. Cell lysates were analyzed by western blotting using an anti-GFP antibody.

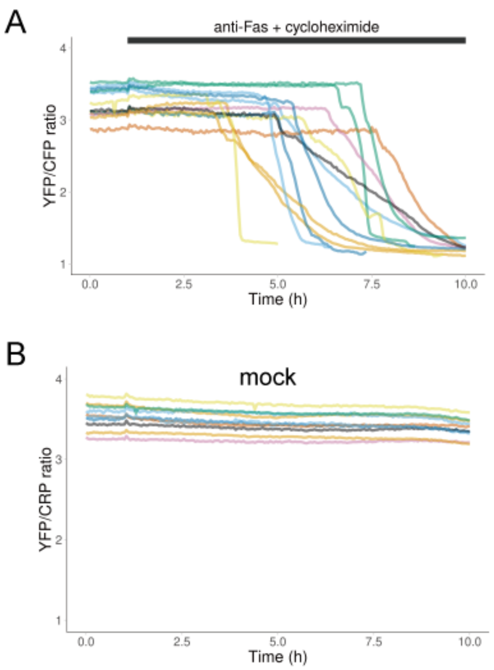


**Figure S2. Dynamics of cytosolic ATP levels in individual apoptotic cells.** Each line represents the time course of the FRET/CFP ratio of AT1.03CR in a single apoptotic cell. Medium with (A) or without (B) anti-FAS and CHX was added at 1 hour after starting imaging. One data set out of three biological replicates are shown (13 cells for anti-FAS treatment and 10 cells for mock treatment).

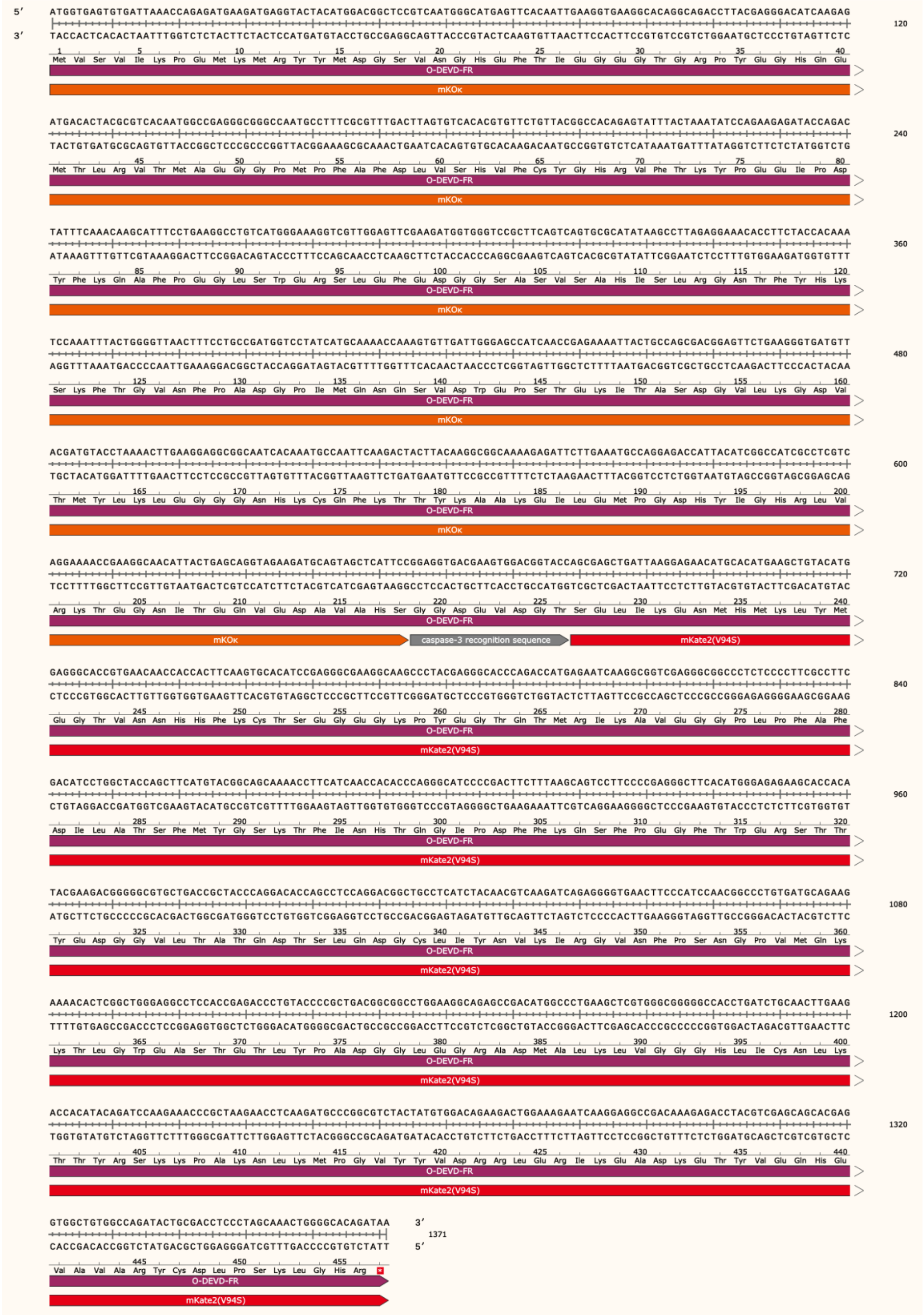


**Figure S3. Nucleotide and amino acid sequences of O-DEVD-FR.**

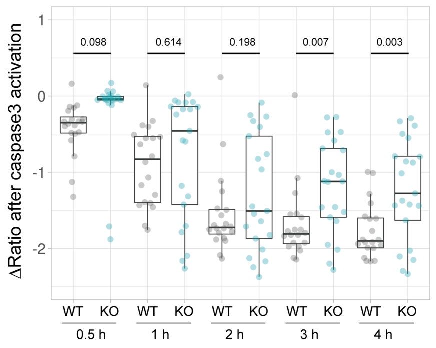


**Figure S4. Knockout of PANX1 suppresses the cytosolic ATP decrease of SW480 cells during apoptosis.**

Apoptosis of wild-type and PANX1-KO SW480 cells expressing AT1.03CR and O-DEVD-FR were induced by TRAIL. Changes in YFP/CFP ratios at indicated time after the onset of caspase-3 activation were calculated for each apoptotic cell (20 [WT] and 21 [PANX1-KO] cells from 3 biological replicates). P-values of Student's t-test were shown.

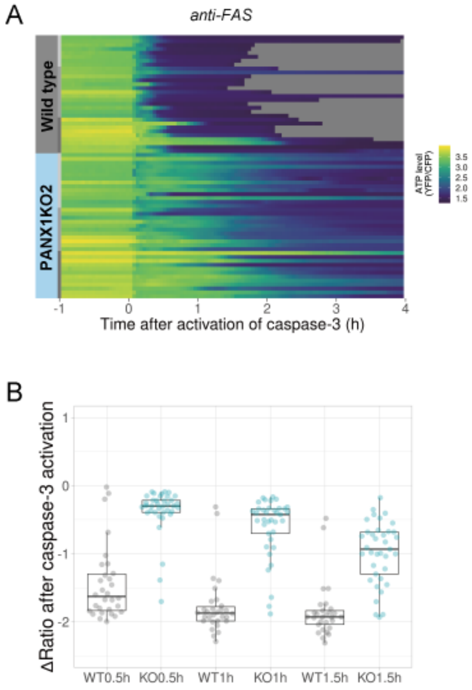


**Figure S5. PANX1 knockout significantly suppresses the decrease in cytosolic ATP levels of apoptotic HeLa cells in an OXPHOS-dependent culture condition.** HeLa cells were cultured in glucose-free DMEM supplemented with 10% FBS, 10 mM lactate and 10 mM DCA. Apoptosis was induced with anti-FAS/CHX. (A) Each line represents the YFP/CFP ratio of AT1.03CR from a single apoptotic cell, and was adjusted by setting the onset of caspase-3 activation as time = 0 (30 [WT] and 37 [PANX1KO2] cells from 3 biological replicates). (B) Effect of PANX1-KO on the decrease in cytosolic ATP levels. Changes in YFP/CFP ratios at indicated time after the onset of caspase-3 activation were calculated for each apoptotic cell. Apoptosis was induced by anti-FAS and cycloheximide.

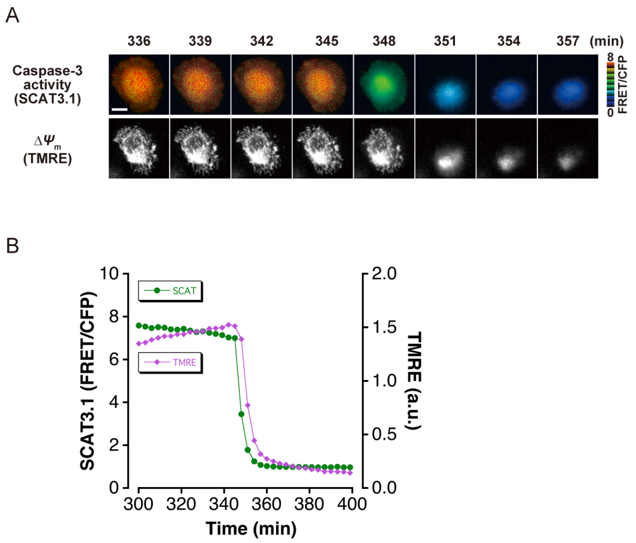


**Figure S6. The initiation of caspase-3 activation occurs almost simultaneously with mitochondrial membrane potential loss.** (A) Time-lapse images of caspase-3 activity and mitochondrial membrane potential ($\Delta\Psi_m$) of a representative single apoptotic cell. Fluorescence images of a HeLa cell, which expressed SCAT3.1 and which was loaded with TMRE, were captured every 3 min. Imaging was initiated 30 min after induction of apoptosis with anti-FAS antibody/CHX. Pseudocolored FRET/CFP ratio images of SCAT3.1 are shown in the upper panel, and TMRE images are shown in the lower panel. Bar, 10 μm. (B) Representative time course of the cytosolic ATP level and $\Delta\Psi_m$ of a single apoptotic cell. Quantified FRET/CFP ratio (green circle, left axis) and TMRE fluorescence intensity (magenta triangle, right axis) from the apoptotic cell in (A) are shown.

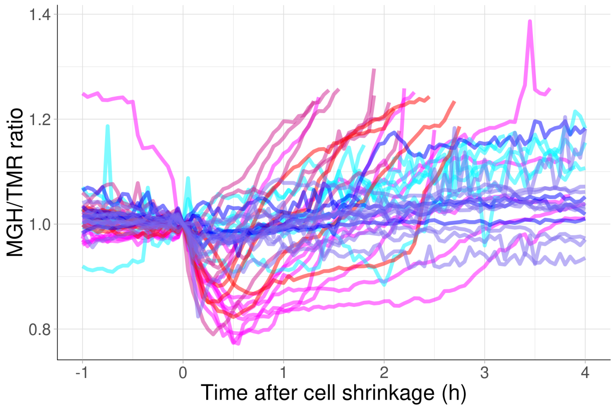


**Figure S7. PANX1 regulates the cytosolic free $Mg^{2+}$ level in apoptotic cells.** HeLa cells expressing Halo-tag were loaded with MGH(AM) and Halo-TMR, followed by induction of apoptosis. Each trace represents a time course of normalized MGH/TMR fluorescence ratios of individual cell from three biological replicates. Timing of the initiation of cell shrinkage was defined as time = 0. Reddish and bluish traces represents wild-type cells (n = 22) and PANX1-KO2 cells (n = 20), respectively, from three biological replicates. Traces from different biological replicates were shown in different color codes.

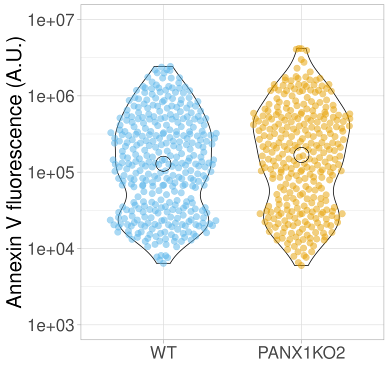


**Figure S8. PANX1 knockout does not affect the externalization of phosphatidylserine.** Apoptosis of HeLa cells were induced by anti-FAS and cycloheximide in the presence of Alexa647-annexinV and propidium iodide. Fluorescence intensity of Alexa647-annexinV of each cell that does not show propidium iodide fluorescence was quantified using a fluorescent microscopy at 6 hours after induction of apoptosis (359 [WT] and 322 [PANX1-KO2] cells from 3 biological replicates).

**Legends for supplementary movies**

**Supplementary movie 1. Time lapse fluorescent movie of an apoptotic wild-type HeLa cell.** YFP (FRET) fluorescence from ATeam is shown. Apoptosis was induced by anti-FAS and cycloheximide. Time = 0 indicates the onset of caspase-3 activation.

**Supplementary movie 2. Time lapse fluorescent movie of an apoptotic PANX1-KO HeLa cell.** The same experimental conditions as supplementary figure 1.

**Supplementary movie3. Time lapse florescent movie of an untreated apoptotic PANX1-KO HeLa cell.** Apoptosis was induced by anti-FAS and cycloheximide. Time = 0 indicates the onset of caspase-3 activation.

**Supplementary movie 4. Time lapse fluorescent movie of a 2DG-treated apoptotic PANX1-KO HeLa cell.** Apoptosis was induced by anti-FAS and cycloheximide. Time = 0 indicates the onset of caspase-3 activation. 2DG was added between 60-63 min.
